# Possible Protective effect of Unani formulation against Isoproterenol induced toxicity in SVEC cells

**DOI:** 10.1101/062802

## Abstract

Main objective of the present study is to screen the protective efficiency of *Khamira Gaozaban Ambari Jadwar Ood Saleeb Wala* (KGAJOSW) a Unani formulation by using an experimentally induced toxicity model. Study mainly focused on the protective efficacy of the Unani formulation “KGAJOSW” with special emphasis to Isoproterenol (ISO) induced toxicity. This formulation has been referred as a “cardio tonic”, both in ancient (Uddin Mk, 1967) and recent literatures (Ahmad et al, 2010) of Unanipathy. The outcome of this study will deliver the protective and non-toxic nature of the KGAJOSW. This will enhance the medicinal properties of other Unani formulation and making them non-toxic.

Existing medicines and therapeutic procedures are target specific and have several post therapeutic complications like hypertension, Obesity, etc. Recent studies demonstrate that non-invasive treatment is the better way to treat and control development of chronic diseases like atherosclerosis. Due to these undesirable effects of synthetic drugs, the world population is turning towards medicines derived from locally available native extracts of edible plant parts and plants products, like Unani and Ayurveda.

In this context it is proposed to investigate the following objectives, which emphasis the lethality, ROS scavenging and expression of SOD-2 protein by KGAJOSW. The outcome of this study will focus on the mode of action and brief mechanisms operative at the “cardio tonic” level as affirmed for KGAJOSW in Unani pharmacopeia.

**Objectives:**

- To identify the lethal dose of Isoproterenol and Unani formulation
- To measure Reactive Oxygen Species (ROS) levels
- To check the Superoxide dismutase2 (SOD2) protein and RNA expression
- Wound healing assay to know the cell proliferation

**Study design:** We come-up with a pilot study to screen the protective as well as non-toxic effects of said Unani formulation. To achieve this we adopted a model of Isoproterenol (ISO) induced toxicity which is widely used to study the myocardial infarction in cells and animals. In this study we want to induce toxicity in simian virus derived murine endothelial cell line (SVEC) cells by treating with ISO and we were assuming this toxicity could be rescued by treating with Khamira (see the Schematic representation). We designed four experimental conditions like Untreated- SVEC cells without any treatment, ISO- SVEC cells receiving Isoproterenol, Khamira- SVEC cells receiving Unani formulation, ISO+Khamira- SVEC cells receiving both Isoproterenol and Khamira.

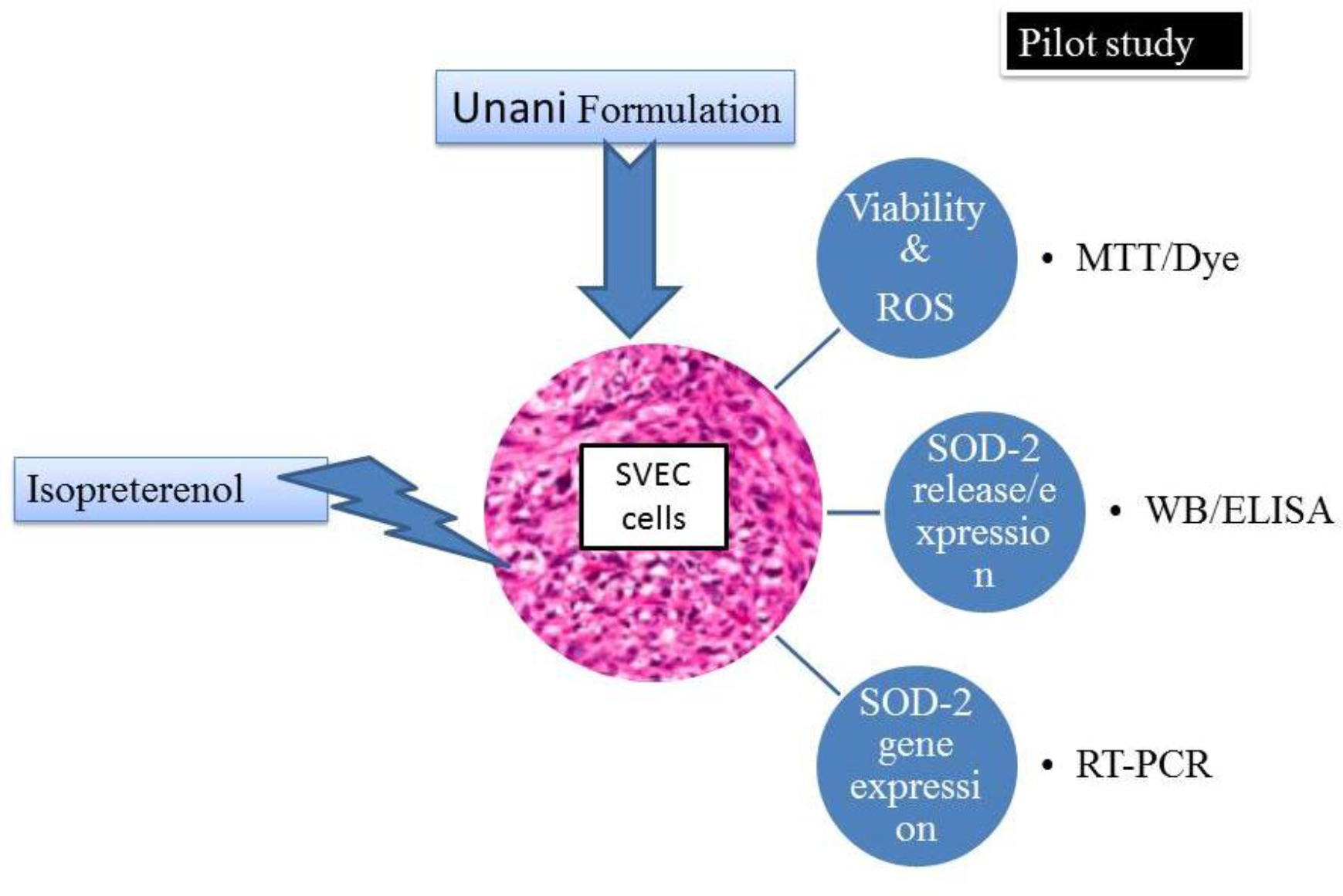

## Materials and Methods

### Materials

SVEC cells, Isoproterenol, KGAJOSW-Unani formulation (Hamdard pharmaceuticals), DMEM, 10%FBS, Western Blotting apparatus, TRIzol reagent, DCF-DA, MTT, Krebs-Ringer Solution [HEPES Buffered] and common chemicals and reagents.

### Cell culture

Cell lines used were: SVEC (from Maneesha Inamdar, MBGU, JNCASR, Bangalore). All cell lines were cultured in DMEM containing 10%FBS (cat. no. 10270, Gibco-BRL, USA) and 2 mM Glutamax (cat. no. 35050, Invitrogen, Carlsbad, USA,).

**Table 1.** Composition of KGAJOSW.

| Ingredients | Quantity |
| --- | --- |
| Gaozaban (*Borago officinalis*) | 30 g |
| Gul-e-Gaozaban (*Borago officinalis*) | 10 g |
| Kishneez khushik (*Coriandrum sativum*) | 10 g |
| Aabresham (*Bombbyx mori*) | 10g |
| Behman surkh (*Salvia haematodes*) | 10 g |
| Behman safaid (*Centaurea behen*) | 10 g |
| Sandal safaid (*Santalum album*) | 10 g |
| Badranjboya (*Mellisa paraviflora*) | 10 g |
| Tukhm-e-Balango (*Lallemantia royleana*) | 10 g |
| Tukm-e-Faranjmushk (*Ocimum gratissimum*) | 10 g |
| Ustukhuddus (*Lavendula stoechas*) | 10 g |
| Toodari Safaid (*cheiranthus cheiri*) | 10 g |
| Toodari Surkh (*Mathiola incana*) | 10 g |
| Amber (*Ambra grasea*) | 2 g |
| Jadwar (*Delphinium denudatum*) | 10 g |
| Ood Saleeb (*Paeonia emodi*) | 10 g |
| Arq-e-Keora (*Pandanus tectorius*) | 10 ml |
| Warq-e-Nuqra (silver) | 6 g |
| Warq-e-Tila (Gold) | 6 g |
| Nabat Safaid | 1 kg |

### Methods

#### MTT assay

MTT ( 3-(4,5-dimethylthiazol-2-yl)-2,5-diphenyltetrazolium bromide) assay, SVEC cells were plated at a density of 500-10,000 cells in 96 well plate and incubated at 37°C, 5%CO2 O/N to allow the cells to attach to wells. ISO, Khamira and ISO+Khamira were added at different concentrations and incubated for 4 h and 24 h. MTT was added to a final concentration of 0.5 mg/ml and incubated at 37°C, 5%CO2 for 3 h to allow MTT to be metabolized. Media was removed and 200 μl of DMSO was added to resuspend formazan (MTT metabolic product) and kept on a shaking table for 5 min. Absorbance was read at 560nm and background at 660 nm was substracted. Absorbance is related with cell quantity. Assay was done in triplicates.

## Measurement of intracellular ROS

DCF-DA is a non-specific probe for ROS to produce fluorescent 2′-7′-dichlorofluorescin. The acetate groups of the DCF-DA are cleaved *in vivo* by indigenous esterases, and subsequent oxidation by intracellular peroxides produces the fluorescent DCF-DA (Thannickal and Fanburg, 2000), which can be detected by flow cytometry. Human ESCs were mechanically detached from culture dishes. Colonies of hESCs were treated with trypsin/EDTA to make single cells. Cells were washed with Krebs-Ringer-HEPES (KRH) buffer containing 1 mM calcium, 0.1 mM glucose, and 0.2%BSA. These cell suspensions were incubated with 10 μM DCF-DA in a 37°C water bath for 30 min. To prevent degradation of DCF-DA from lights, we performed the reaction in the darkness.

## Krebs-Ringer Solution [HEPES Buffered]

120 mM NaCl, 5 mM KCl, 2 mM CaCl2, 1 mM MgCl2, 25 mM sodium bicarbonate, 5.5 mM HEPES, 1 mM D-glucose, pH 7.2 ± 0.15. +0.2%BSA

**Table: 2.** Krebs-Ringer Modified Buffer (KRB) Adjust the pH to 7.4. Most experiments are performed in buffer supplemented with 1 mM CaCl_2_ and add 0.2%BSA

| Reagent | Final concentration |
| --- | --- |
| NaCl | 135 mM |
| KCl | 5 m M |
| MgSO<sub>4</sub> | 1 m M |
| K<sub>2</sub>HPO<sub>4</sub> | 0.4 mM |
| Glucose | 5.5 m M |
| HEPES | 20 m M |

### Wound healing Assay

SVECs were grown to 100%confluence till the cell monolayer was wounded using a 200 μl pipette tip, gently washed with PBS and incubated with fresh media supplemented with 5%FBS and 1,5 and 10 mg/mL Conc. of Khamira respectively. Cells were monitored using IX70 inverted fluorescence microscope (Olympus, Tokyo, Japan) and imaged at different time intervals using a CCD camera (CoolSNAP; Roper Scientific, Inc, Trenton, NJ). The rate of wound closure was measured by calculating the distance covered.

### Western Blotting

For protein lysates, cells were pelleted, washed with PBS and lysed in a buffer containing 25mM HEPES, 150mMNaCl, 1%NP-40, 10%glycerol, 10mM MgCl2, 1mM EDTA, 1mM Sodium ortho vanadate and 10 μg/ml protease inhibitor cocktail (SIGMA Chemical Co., USA)on ice and clarified by centrifugation. Protein was estimated using Bradford reagent and 50 μg of lysate was loaded on 12%SDS-PAGE. Gel was electroblotted on PVDF membrane, probed with specific antibody and developed using ECL chemiluminescence (Biorad,US).

### Total RNA isolation

Total RNA was isolated from SVEC cell lines using TRI reagent following the manufacturer’s protocol (Sigma). 1–2 × 106 cells were dislodged from culture dish, washed with PBS and lysed in 1 ml of TRI reagent and allowed to stand for 5 min at room temperature (RT). 200 μl of distilled chloroform was added, mixed vigorously by vortexing and kept at RT for 15 min. The resulting mixture was centrifuged at 12,000 × g (Sigma 3K30) for 15 min at 4°C. The upper aqueous phase was carefully transferred to another tube and RNA was precipitated by the addition of 500 μl isopropanol followed by thorough mixing and incubation at RT for 10 min. RNA was pelleted by centrifuging at 12,000 × g for 10min at 4°C and washed twice with 75%ethanol. The RNA pellet obtained was allowed to air dry to remove traces of ethanol and resuspended in 30 μl of DEPC treated water. The concentration of RNA was determined by using Nanodrop by taking absorbance of 1:500 dilution of the RNA at 260 nm and calculated by using the formula: Concentration of RNA in μg/ml = A260 × 40 × dilution factor. RNA preparation was checked on 1.0%agarose gel.

### First strand cDNA synthesis

1 μg of RNA was used for cDNA preparation using First strand cDNA synthesis kit according to manufacturer’s instructions (Fermentas). The reaction mix (Reaction mix-1) containing 1 μg of RNA and oligo dT was made upto 11 μl with DEPC water. Reaction was prepared on ice, mixed gently and incubated at 70 °C for 5 min. After incubation, other components (Reaction mix-2) i.e 5X reaction buffer, RNAse out RNAse inhibitor and dNTP mix were added. The components were mixed gently and incubated at 42°C for 2 min. 3 μl reverse transcriptase enzyme was added to make up the reaction volume to 20 μl. The reaction was incubated at 42°C for 50 minutes, 70°C for 15 minutes followed quick chill and add 30 μl of nuclease free water.

**Table 3:** Reaction mix-2. *Reaction mix 1: 3 μg of RNA, Oligo dT 1 μl, Nuclease free water = total vol 8 μl

| Reaction mix | Volume |
| --- | --- |
| Reaction mix 1 | 8.0 $\mu\text{l}$ |
| 5X reaction buffer | 4.0 $\mu\text{l}$ |
| RNaseout | 0.5 $\mu\text{l}$ |
| dNTP mix (2 mM) | 5.0 $\mu\text{l}$ |
| Total | 17.0 $\mu\text{l}$ |
| Reverse Transcriptase (Superscript) | 3.0 $\mu\text{l}$ |
| Total reaction mix | 20.0 $\mu\text{l}$ |

\*Reaction mix 1: 3 $\mu\text{g}$ of RNA, Oligo dT 1 $\mu\text{l}$ , Nuclease free water = total vol 8 $\mu\text{l}$

### Quantitative Real-Time RT-PCR Assay

Real-time RT-PCR reactions were performed using Real Time PCR Detection System (Light Cycler System, Roche Diagnostics, Germany) with a mixture composed of SYBR Green PCR Master Mix, primers and cDNA. The reactions were carried out for 40 cycles at 95°C denaturation for 10 sec, 55°C annealing for 30 sec and 72°C elongation for 30 sec. The primer sequences were as follows: SOD-2, forward primer-5’-CAGACCTGCCTTACGACTATGG-3’ and reverse primer-5’-CTCGGTGGCGTTGAGATTGTT-3’; GAPDH, forward primer −5’-AGGTCGGTGTGAACGGATTTG-3’ and reverse primer-5’-TGTAGACCATGTAGTTGAGGTCA-3’. The copy numbers of SOD-2 mRNA were quantified by determining the Cp values, followed by normalization to GAPDH.

## Results and Discussion

### Cell viability Assay

To check for cell viability, SVEC cells were treated with ISO and Khamira at different concentrations individually. Cells were treated with different concentrations of ISO i.e., 100 μM, 150 μM, 200 μM and 250 μM for 24 h. Similarly another set of cells were treated with different concentration of Khamira i.e., 1, 10, 20 and 40 mg/mL for 24 h. Later MTT assay was performed to observe the concentration at which ISO and Khamira were toxic for the cells. Earlier studies have reported toxicity of ISO at 180 mg/Kg body weight in rat models. Till to date nobody reported any significant effect of Khamira on Cell lines. Hence in this study we made an attempt to see the effect of Khamira on SVEC cells.

Result shows that ISO at 200 μM and 250μpM shows severe mortality (Figure 1.A) when compared to untreated cells. Khamira at 40 mg/mL shows high mortality (Figure 1.B) rate when compared to other concentrations. Based on these results we have chosen to use 150 μM of ISO and 5 and 10 mg/mL of Khamira for further studies.

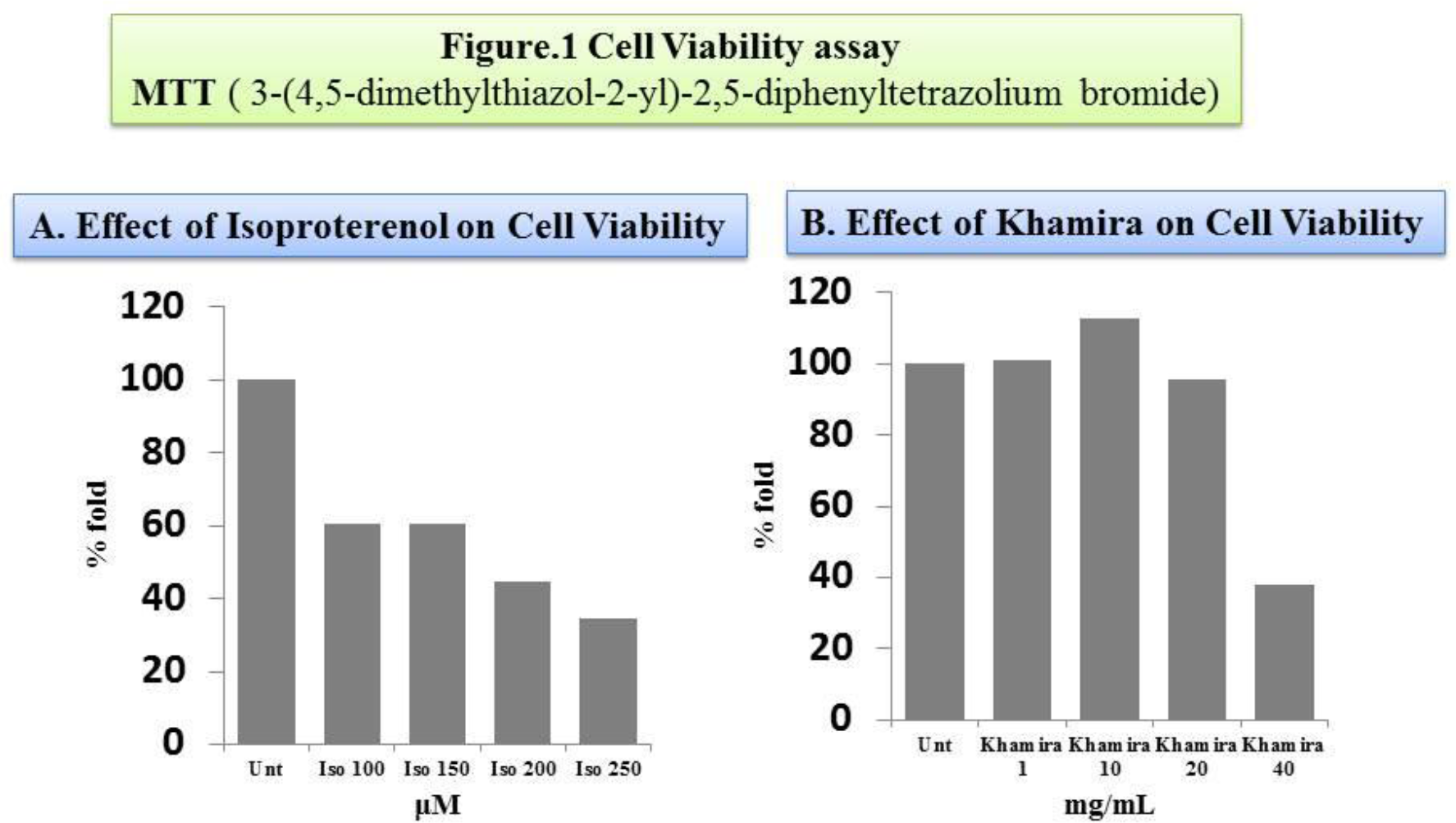

### ROS assay

ROS generated by mitochondrial respiration play an important role in maintaining cellular functions. Increased levels of ROS develops an imbalance between oxidnat and anitoxidnat status that furhter leads to xoidative stress. The production of ROS as natural by products of metabolism is a consequence of using oxygen as an electron acceptor, and when an imbalance in the redox homeostasis occurs, ROS are a considerable cause of DNA damage. The major elements casuing oxidative stress include ROS. We there fore characterized ROS production in the ISO and Khamira treated cells. ISO treatment significantly stimulated ROS production, which was abated by Khamira (Fig.2A. - D), suggesting that Khamira played a protective role via the inhibition of ROS production. The FITC-A vlaues for Untreated, ISO, Khamira and ISO+Khamira are like 59,468; 67,754; 60,872; and 63,984 respectively. We assume that the biochemical basis behind the decreased ROS levels in Khamira treatement could be through increasing endogenous antioxidant levels.

The source of ROS is the key determinant of its subsequent effects. Collectively, the present study showed that Khamira ameliorated pathological alternations via specific inhibition of ROS production. However, the relationships between antiooxidant enzymes and Khamira are still unknown. It has been speculated that Khamira may reduce the lipid levels or cardiotonic specific to strenthening the heart muscles, to inhibit cardiacpathology, which is supported by a our study. However, the mechanism remains to be further elucidated.

**Figure 2A.**
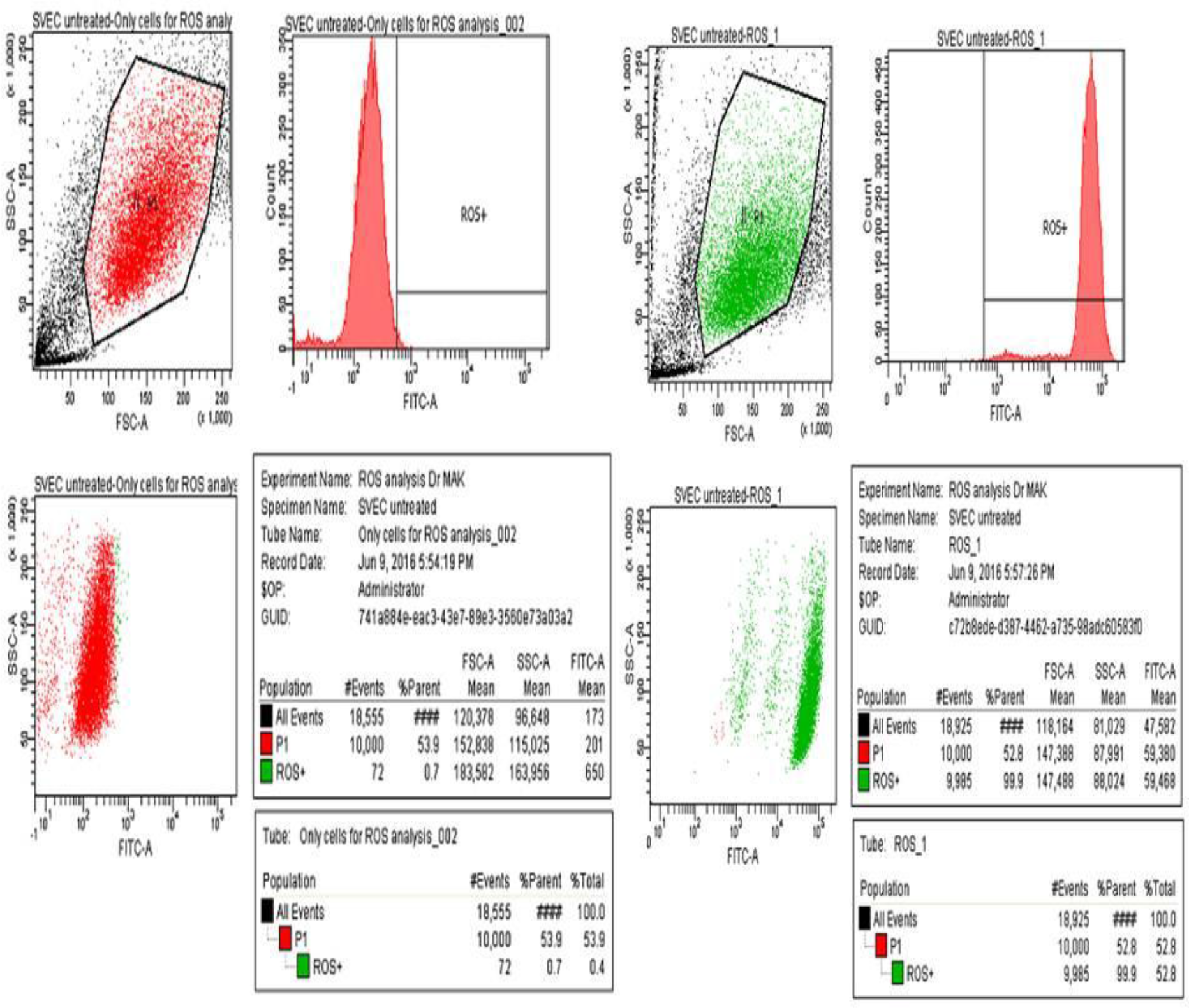
Expressin of ROS in Untreated SVEC cells.

**Figure. 2.B.**
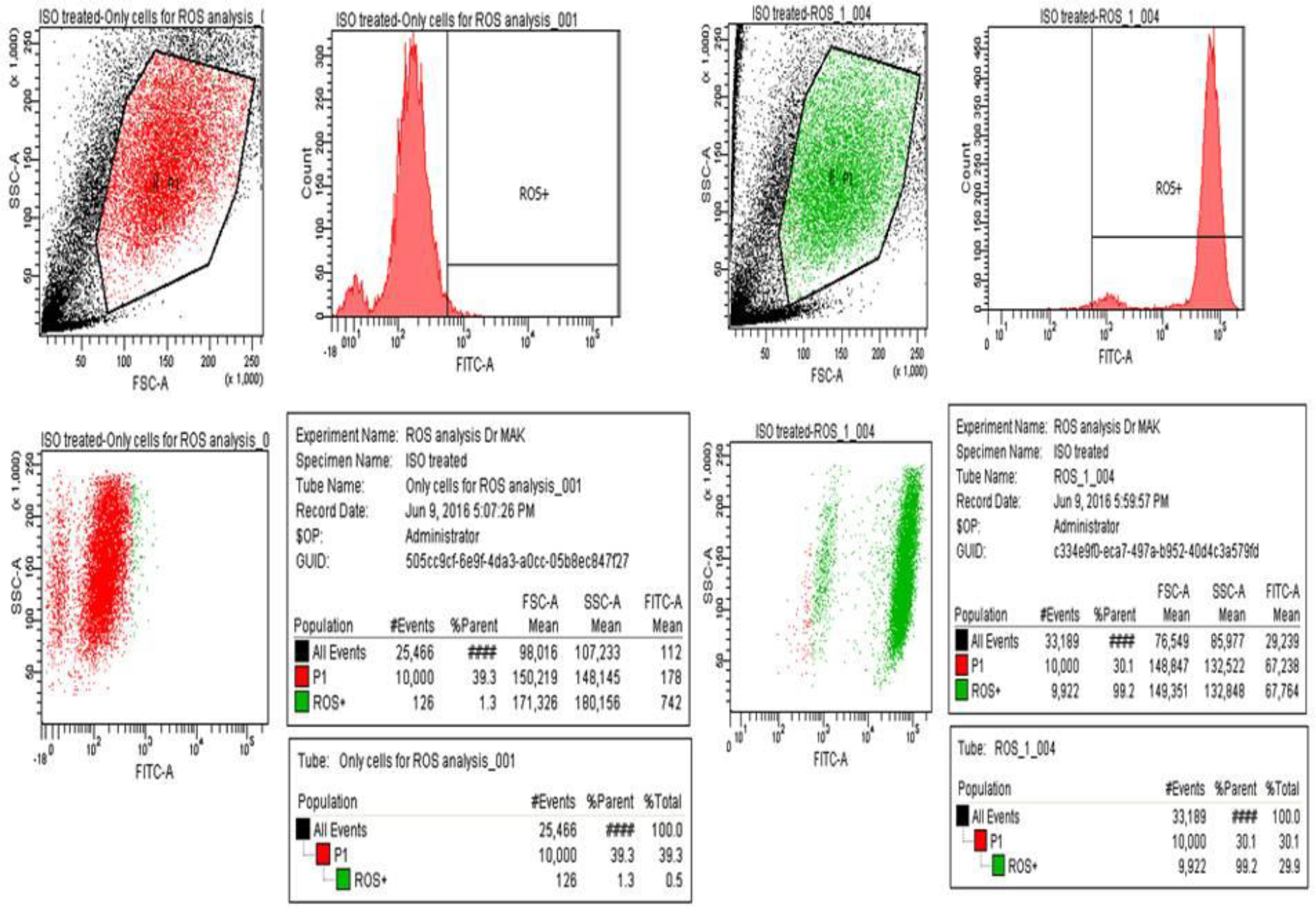
Expession of Ros in ISO treared SVEC Cells.

**Figure. 2.C.**
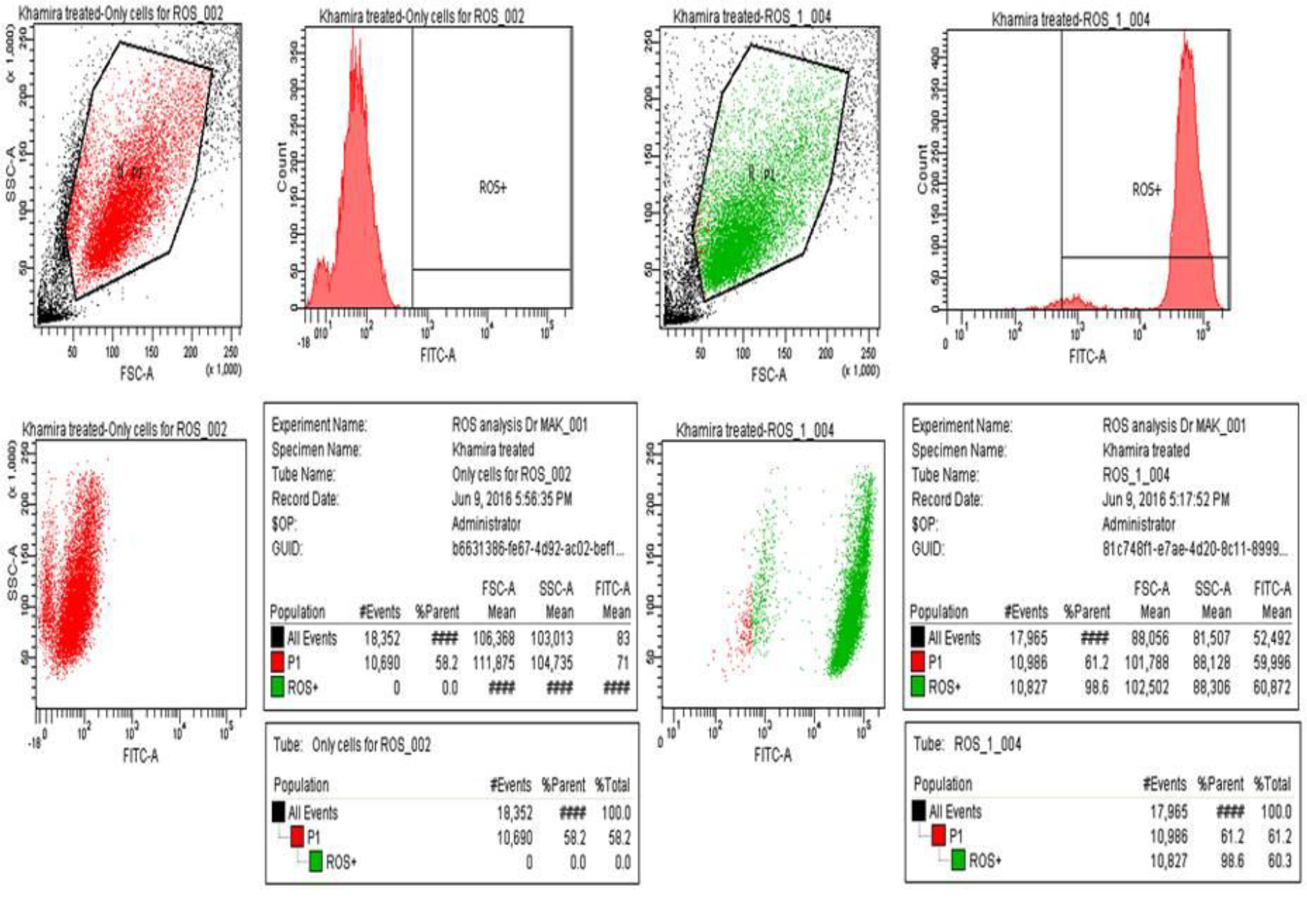
Expression Of ROS in Khamira treated SVEC cells.

**Figure. 2.D.**
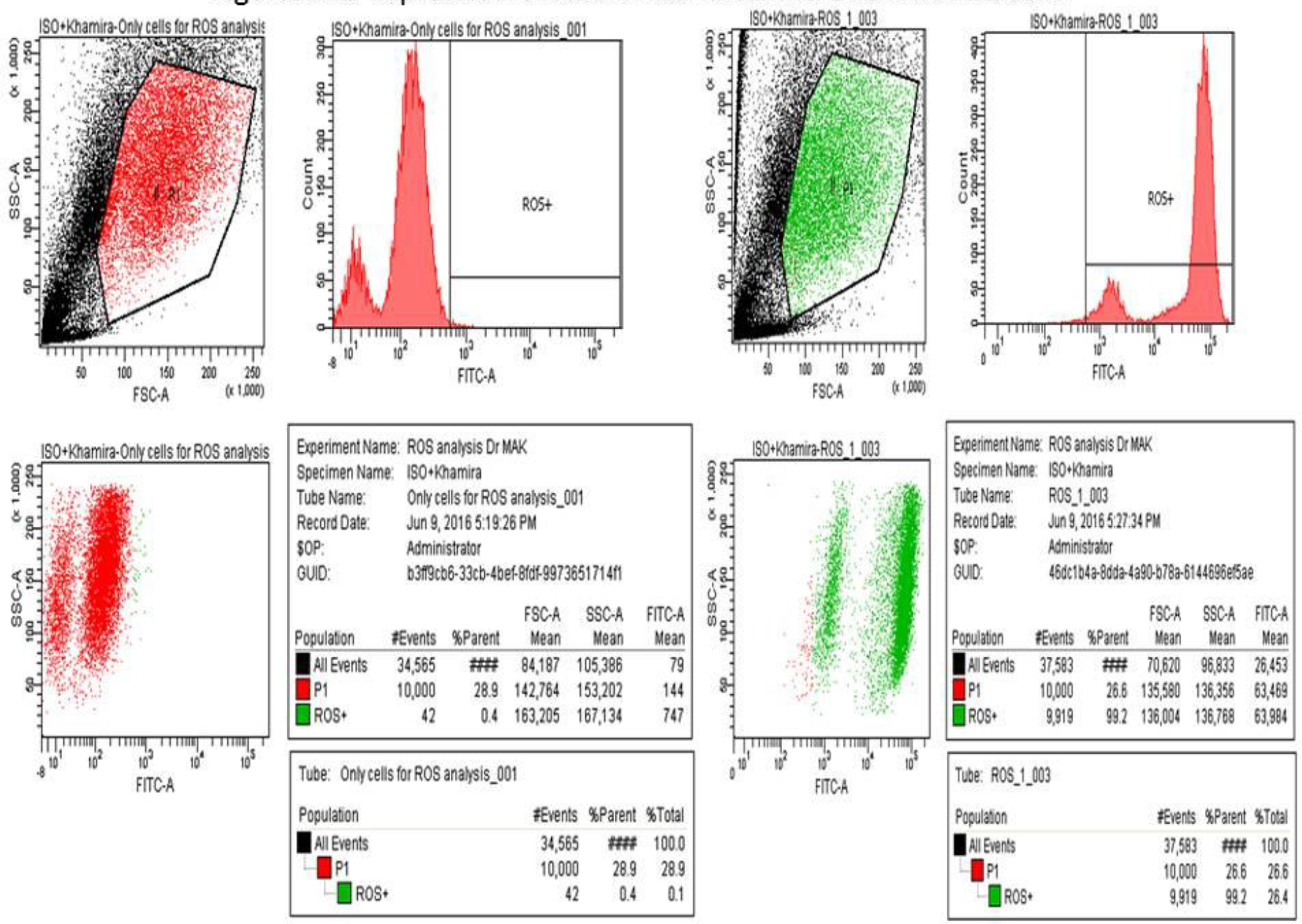
Expression of ROS in ISO+ treated SvEC cells.

### Wound healing assay

The wound-healing assay is simple, inexpensive, and one of the earliest developed methods to study directional cell migration in vitro. This method mimics cell migration during wound healing in vivo. The basic steps involve creating a “wound” in a cell monolayer, capturing the images at the beginning and at regular intervals during cell migration to close the wound, and comparing the images to quantify the migration rate of the cells.

Wound healing assay was performed to check if Khamira induced cell proliferation. SVEC cells treated with Khamira showed rapid increase in cell proliferation rates to cover the wound when compared to control cells which migrated at a slower rate to close the wound (Figure 3A &3B). Fig.3.C shows that Khamira with 5 mg/mL shows a rapid wound healing ability than the other concentrations. MTT assay and Wound healing assay showed increased cell proliferation and migration in Khamira treatment. The biochemical signalling behind this phenomenon is yet to be explored. However, it might be due to a yet to be known defence mechanism to overcome the toxicity or the ability to heal the wound rapidly.

**Figure. 3A.**
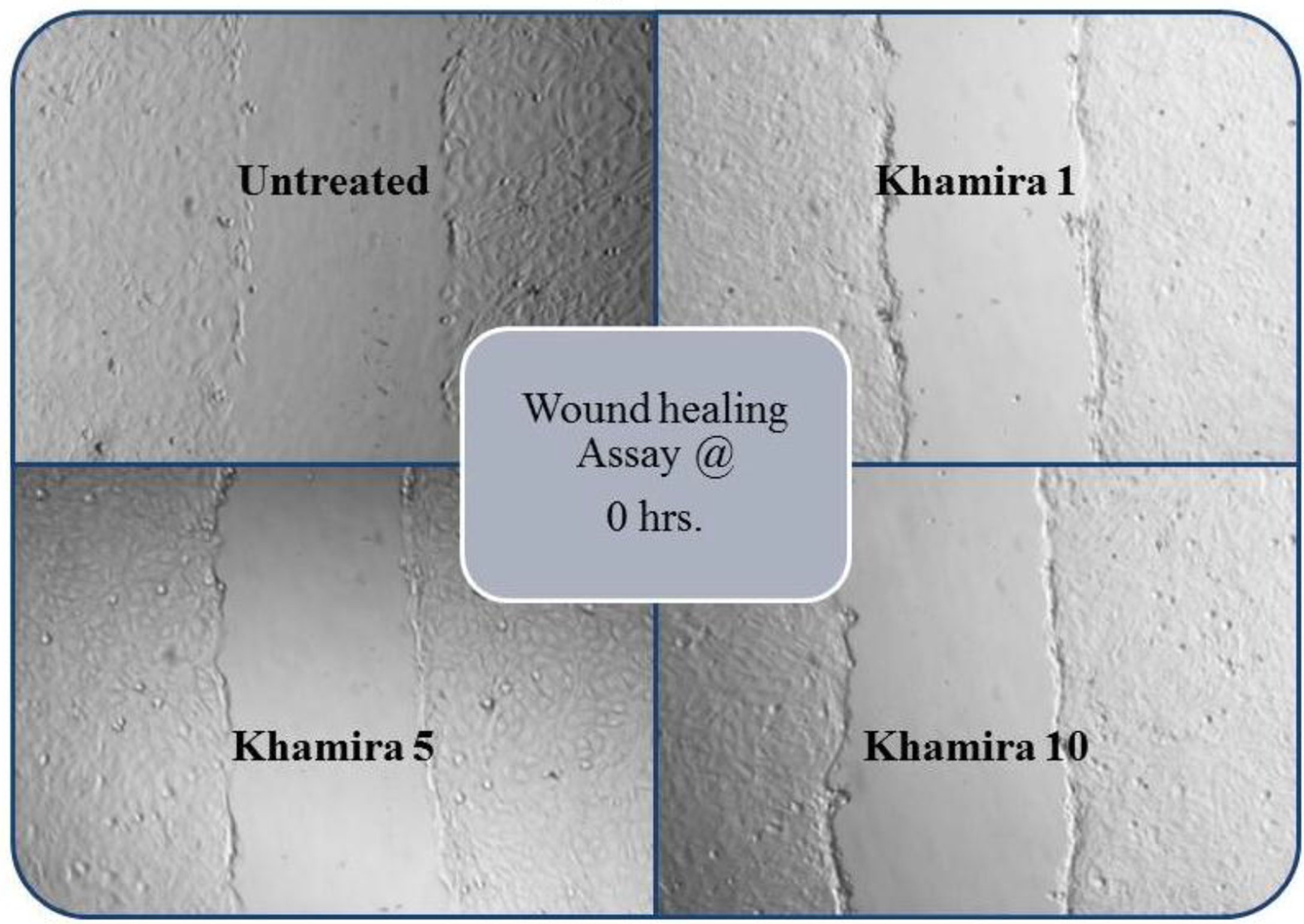

**Figure. 3B.**
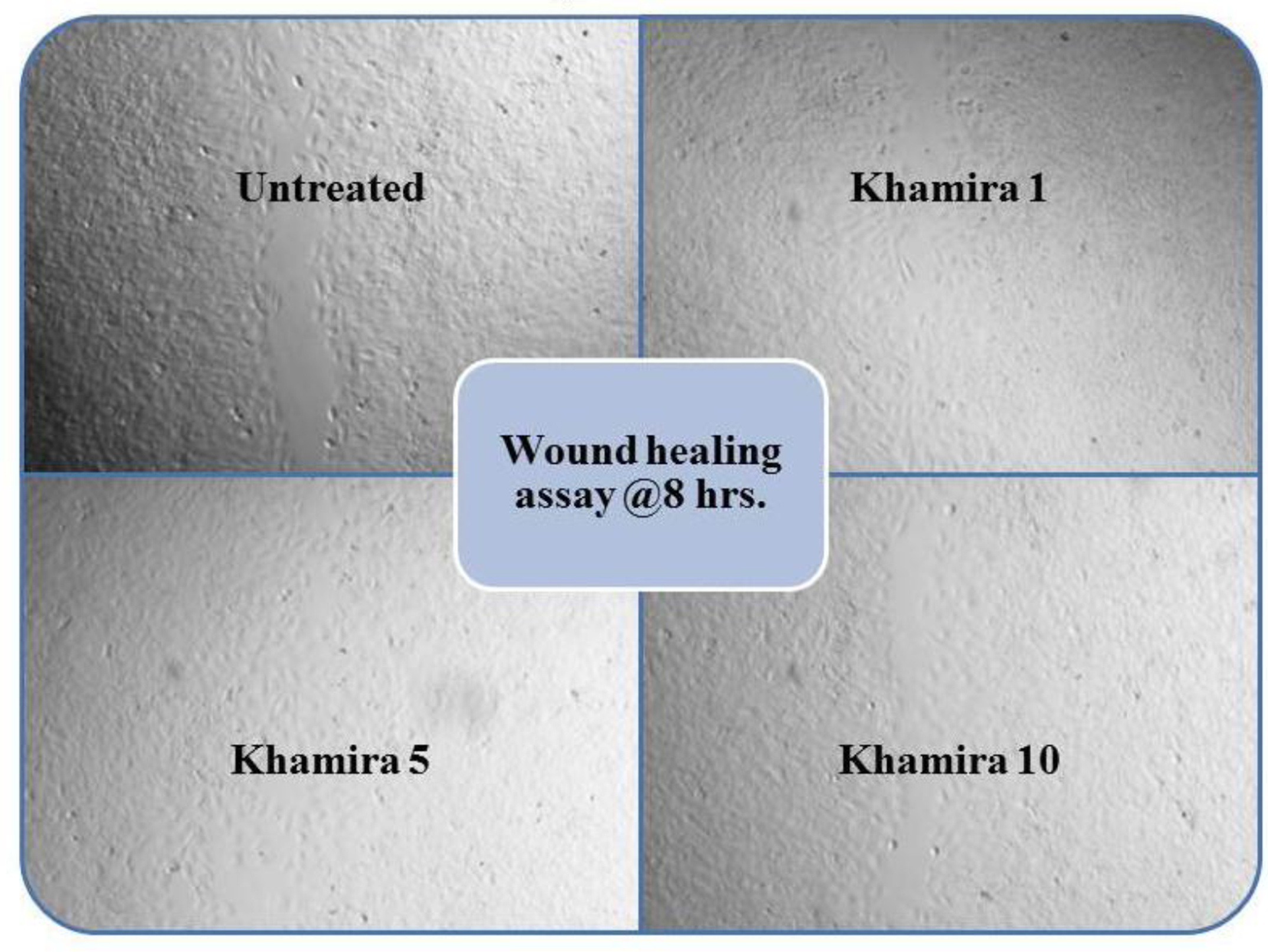

**Figure 3.**
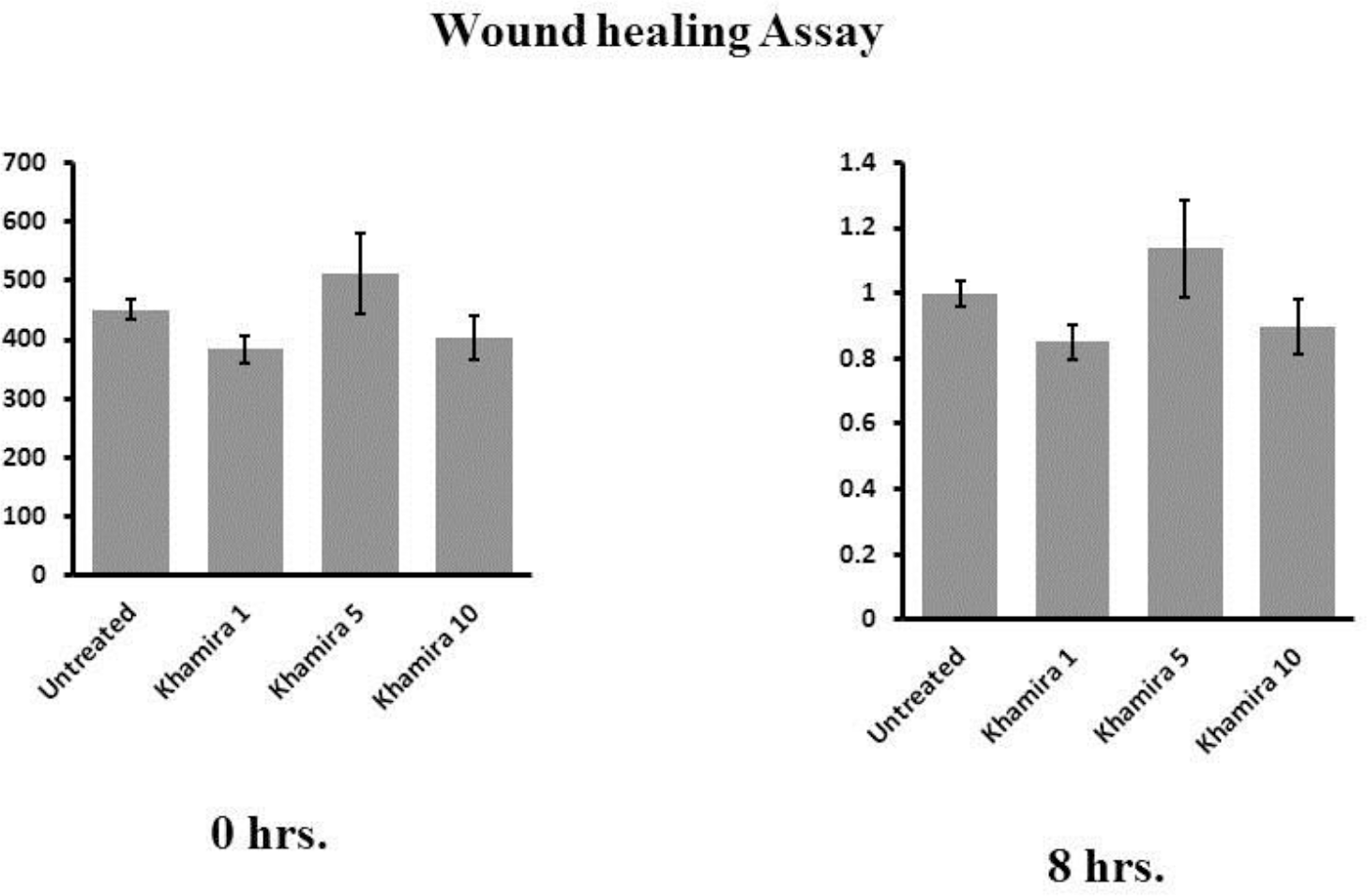
q-PCR Products.

### Protein Expression studies

Taking the cues from ROS and MTT we further moved to know the protein expression under the proposed experimental conditions. If this context we tried to explore the expression of SOD2 a potent antioxidant enzyme of mitochondrial origin. By doing protein estimation by Bradford’s and Coomassie staining (Fig. 4) we were able to know the concentrations and presence several proteins in all the experimental conditions and volume of sample to be loaded.

We tried to check the expression of SOD2 but remain unsuccessful (able to see faint band after long exposure) and able to see the expression of GAPDH at 35 kDa. Literature survey says that expression of SOD2 varies in between 19 to 25 kDa based on its isotype, hence we believe that expression of SOD2 need to be standardized.

**Figure 4.**
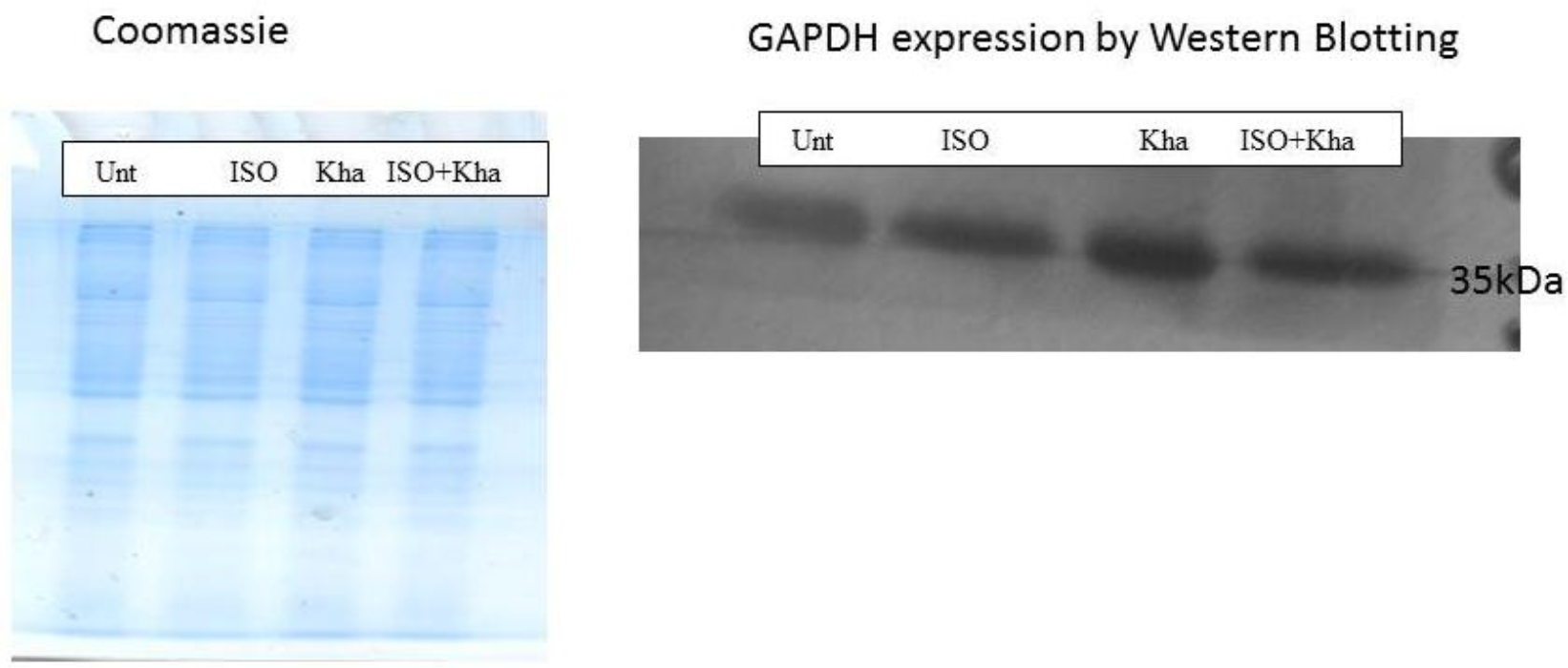
Protein expression.

### SOD2 gene expression by using q-PCR

SOD2 is the member of superoxide dismutases family and plays a vital role in maintaining the homeostasis between ROS and antioxidant status of the cell. After conducting protein expression studies we proceeded further to look into the expression of SOD2 at its genetic level by using q-PCR. To perform this we isolated RNA and constructed c-DNAs for all the fours experimental samples. SOD2 is a mitochondrial enzyme that converts superoxide anion into peroxide, which is substrate for Prdxs and Gpxs that produce H2O as a final product.

Quantitative Real time PCR amplification of DNA was carried out using FastStart SYBR Green Master (Biorad) and specific oligonucleotides in Opticon Real Time DNA engine (Bio-Rad). A melting curve analysis was performed immediately after amplification at a linear temperature transition rate of 0.2°C/s from 61°C to 91°C with continuous fluorescence acquisition. Gene expression was normalized to the housekeeping gene GAPDH. The amplicon size was confirmed by gel electrophoresis.

Levels of SOD2 gene expression in both untreated and Khamira close to each other indicating there is increase in SOD2 expression. Where levels of SOD2 in both ISO alone and ISO+Khamira treated cells levels are increased to 1.8 and 2.08 indicating there is some ROS generation which is inducing SOD2 expression. When we compared SOD2 data with ROS analysis we found that, ROS levels are also increased in ISO and ISO+Khamira cells.

**Figure 5.**
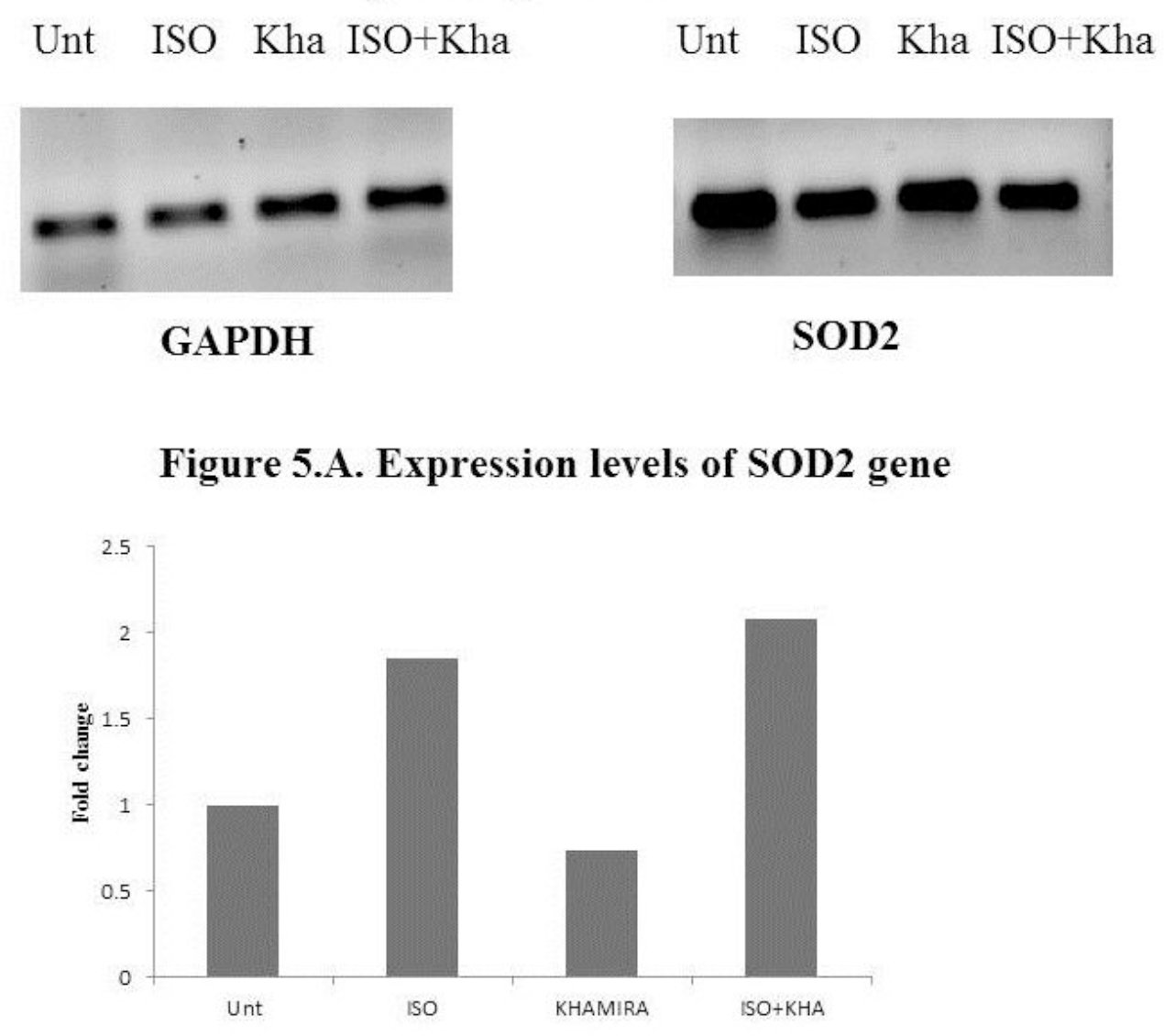
q-PCR Products.

## Conclusion

This pilot study shows that Khamira is nontoxic to SVEC cells up to a concentration of 20 mg/mL after this concentration cells are unable to survive. We also observed that the 150 μM concentration of Isoproterenol induces mild lethality and anything above this concentration leading to increased toxicity and induces death. During this study we found rapid filling of cells in wounded region of Khamira with 5mg/mL concentration than that of 10 mg/mL. In this entire study we adopted a method of Khamira cotreatment but not pre-treatment. When we look into the ROS values we assume that Khamira is able to maintain the sufficient endogenous antioxidant status that is useful to rescue cells from ROS or free radicals. But when we observe the levels of SOD2 it is evident that Khamira might be acting either on signalling events prior to ROS or after ROS generation. The reason for this statement is SOD2 levels increase when there is elevated ROS generation. However, this is a pilot study to look into the possible protective role of Khamira in experimentally induced toxic conditions. More controlled *in vivo* and *in vitro* studies are required to prove the exact mechanism lying behind increased cell viability and wound healing assay.

